# CSMA: an ImageJ Plugin for the Analysis of Wound Healing Assays

**DOI:** 10.1101/2025.01.12.632613

**Authors:** Tri T. Pham, Amina Sagymbayeva, Timur Elebessov, Zhadyra Onzhanova, Ferdinand Molnár

## Abstract

Accurate quantification of wound closure in cell migration assays is crucial yet challenging. Still, existing methods often underperform due to omitting cell detection within the wound area, resulting in biased outcomes. To overcome this limitation, we developed the CSMA plugin for ImageJ. CSMA utilizes advanced image processing techniques, including contrast enhancement, edge detection, and morphological operations, to precisely identify and quantify cells in the wound region. The plugin offers user-friendly features and adjustable parameters to accommodate different imaging conditions, ensuring robust performance across diverse experimental setups. Validation against conventional tools confirms CSMA’s superior ability to delineate wound boundaries and provide accurate estimations of area and width at every time point. As applied to SW480-ADH colon cancer cells treated with various compounds, CSMA proves valuable in biomedical research. It represents a significant advancement in wound healing assay analysis, providing researchers with a simple and reliable tool for studying cell migration dynamics with enhanced precision and reproducibility. CSMA is available as an ImageJ plugin and source code at https://github.com/AminaSagymbayeva/CSMA_WoundHealing.

## INTRODUCTION

Wound healing assay is a classic method for the assessment of collective cell movement in epithelial and endothelial tissue cultures ^1^. It can be applied to study persistence, speed, and the effect of cell-to-cell and cell-to-extracellular matrix interactions in populations of migrating cells ^1–3^. Moreover, it may be coupled with different microscopy methods to study intracellular events that occur during migration ^1^. Wound healing assay is often a method of choice because of its compatibility with multiple cell lines, easy and inexpensive setup, and wide application ^10^. Therefore, it is used as a model for studying many processes, such as embryonic morphogenesis, angiogenesis, *in vivo* wound healing, and metastasis ^4–8^.

Migrating cell cohorts maintain their intercellular junctions, unlike individual migrating cells ^6^. Cell-cell junctions are supported by adherens junction proteins that interact with actin cytoskeleton and allow cell plasticity ^6^. In epithelial cells, cadherins, specifically E-cadherin (*CDH1*), play a central role in cell-cell adhesion and supracellular polarization ^3,6^. Cell polarization is induced by extracellular chemical signals, such as growth factors, cytokines, and extracellular ligands ^3,6^. In response to chemical stimuli, the actin cytoskeletons in cells at the leading edge reorganize to form various protrusions, such as broad lamellipodia and cylindrical filopodia ^3,6^. Interestingly, not only do the cells at the front of the edge develop protrusions, but also those behind them form cryptic lamellipodia to interact with the substratum and thus promote further movement ^9^. Four small GTPases Rho, Rac, Ras, and Cdc42 are the dominant regulators of the collective cell movement, as they control many processes from cell polarization to adhesion, to pseudopodia development ^10^.

Wound healing assay includes growing cells of interest to high-level confluency, creating a narrow gap in the tissue monolayer, and taking snapshots of it at equal time intervals to observe the gap closure ^1^. There are two methods of gap introduction: physical exclusion and direct manipulation ^2^. The first method prevents monolayer growth under an insert placed in the dish. Although it is more expensive, this method causes less damage to cells at the gap front and produces a cleaner gap with less debris ^2^. The second method, commonly referred to as the scratch assay, is more popular due to its cost efficiency and ease of implementation. A gap is created by scratching the surface of the tissue monolayer with a sharp object, such as a micropipette tip ^1^. Subsequently, a time-lapse series of images is captured to monitor and estimate the rate of gap closure ^2^.

Conventionally, each snapshot image is analyzed manually ^2^. However, this method is very time-consuming and prone to user bias. Several plugins, scripts, and stand-alone software have been developed for the automatic or semi-automatic quantification of wound closure. Many of them demonstrate highly accurate wound detection on images with different illumination conditions and require the adjustment of only a few parameters ^11–18^. However, since some of these algorithms only account for the largest cell-free area, as demonstrated in Fig. 1, they sometimes underestimate the true wound area, especially when it is divided into multiple regions. In addition, they detect only the wound front and ignore cells that are left in the center of the gap, which also introduces a bias in the results as shown in Fig. S1. Though not all cell types tend to break away from the tissue monolayer and migrate inside the gap, for those cell lines that do, the currently available software may produce inaccurate results.

**Figure 1:**
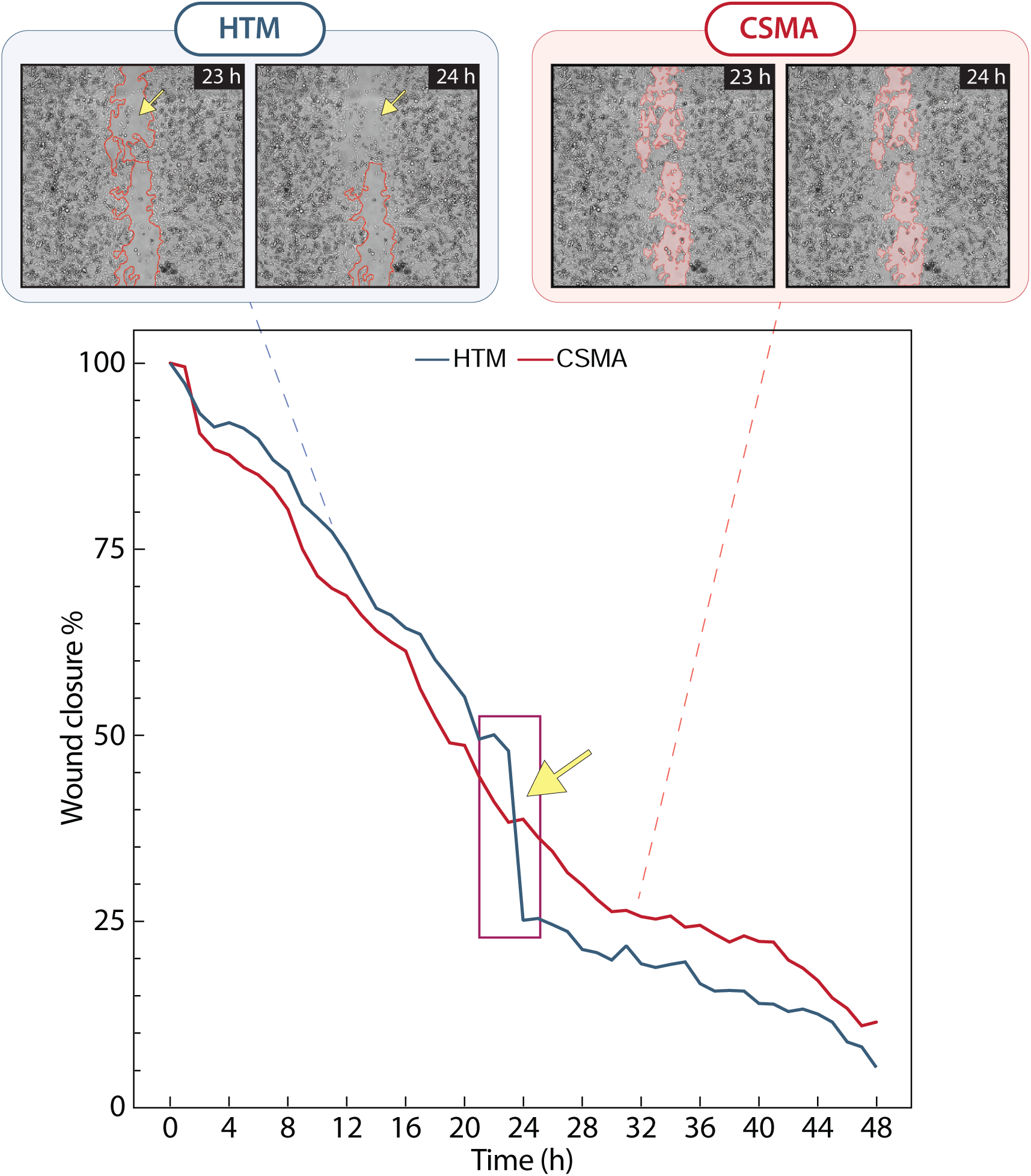
Representative images of wound closure in the solvent control group of SW480-ADH colon cancer cells detected by the High-Throughput Microscopy (HTM) algorhithm and the Cell Scratch Migration Analysis (CSMA) plugin (quantified with default settings). Yellow arrows point to the region dismissed from the calculations due to the closure of the wound gap in the middle. In the corresponding graph, the yellow arrow points to the rapid drop of wound area % as detected with HTM due to the artifact described above.

In this paper, we present a newly developed ImageJ plugin named Cell Scratch Migration Analysis (CSMA), which improves on the existing tools by (i) enhancing the quantification of the wound closure by detecting the cells found in the middle of the wound, (ii) providing a user-friendly interface, (iii) offering two modes of quantification by wound area closure (hence wound closure from this point on) or by wound width, and (iv) supporting various imaging conditions due to the flexibility in parameter adjustment. We demonstrated the features of the applied CSMA plugin on a dataset created by treating SW480-ADH colon cancer cells with weak (WVA) and strong (SVA) vitamin D receptor (VDR) agonists, where WVA is about 10,000 times weaker than SVA. VDR signaling is an excellent model for studying migration because *CDH1*, involved in cell-cell adhesion, is its direct target gene ^19^. CSMA is available as an ImageJ plugin and source code at https://github.com/AminaSagymbayeva/CSMA_WoundHealing.

## MATERIALS AND METHODS

### Cell culture and maintenance

The SW480-ADH human colon cancer cell line, which was used to test the CSMA ImageJ plugin, was kindly provided by Professor Alberto Muñoz Terol from CSIC-Autonomous University of Madrid. The cells were cultured in Dulbeccos Modified Eagle Medium (DMEM) (high glucose, GlutaMAX, Life Technologies Limited, UK) supplemented with 10% (v/v) fetal bovine serum (FBS, Life Technologies Limited, UK), and 1% (v/v) penicillin-streptomycin (#15140-122, Gibco) at 37žC in 5% CO_2_. Prior to experiments, the cells were cultured for three days, reaching 70% confluency. Cell viability was determined by trypan blue (cat. no. T10282, Life Technologies Limited, UK) exclusion using Corning hemocytometer Cell Counter (cat no. CLS6749, Corning, Inc, Germany).

### Wound healing assay

SW480-ADH human colon cancer cells were seeded in a standard 24-well culture plate at a density of 2.5 *×* 10^5^ cells per well and incubated in 500*µl* of DMEM supplemented with 10% (v/v) FBS at 37žC until they reached the confluency of 90 - 95%. All experiments were performed on cells between passages 4 and 10. A gap in the cell monolayer was introduced using a sterile generic 200*µl* micropipette tip. The 200*µl* micropipette with a plastic tip was pressed against a plastic ruler pre-sterilized with 70% ethanol solution to make the gap in a straight line at the middle of the well. Subsequently, the growth medium was removed, and the well was gently washed two times with 1*×* phosphate buffer solution (PBS) to remove detached cells and cell debris. Cells were treated with DMSO as a solvent control, 10*nM* SVA, or 100*µM* WVA. Prior to the treatment, both ligands were dissolved in 500*µl* DMEM supplemented with 5% FBS. The cells were then incubated at 37žC in 5% CO_2_ for 48 hours. The experiment was performed once with four replicates per condition.

### Datasets for testing and analysis

The images were acquired using an Omni Cell Imager Microscope (CytoSMARTő Technologies BV, The Netherlands) at 10*×* magnification. The imaging was done at 1-hour intervals over 48 hours. Regions with 1 mm radii were selected from each well at approximately the same position. Regions close to the well periphery were avoided during selection due to poor illumination. A total of 49 images were exported for each condition as JPEG files. Regions with dimensions of approximately 1537 *×* 1537 pixels were cropped from the exported files using the ImageJ toolkit.

Additionally, a publicly available dataset, consisting of 48 images of mutant human renal carcinoma cell line, 769-P (ATCC CRL-1933) imaged for 23.5 hours every 30 minutes using the Live Cell-R Station (Olympus), was used for the comparative testing of different wound detection tools ^14^.

### Data analysis with CSMA

The CSMA ImageJ plugin is free and open-source software based on the OpenCV (version 4.7.0.72) library. The plugin was developed in both Python 3 and Java programming languages. Python 3 is used for image processing and the user interface (UI), whereas Java facilitates communication between ImageJ and Python.

The image analysis tool is available as the ImageJ plugin for Windows and a source code with UI for Linux and macOS. The plugin supports common image extensions such as *.tif, *.jpeg, and *.png. Before using the plugin, the user must configure a virtual environment named ImageJCSMA using an executable file included in the package. All necessary Python libraries are automatically installed in this environment, preventing conflicts with possibly existing library versions on the users PC.

The image processing algorithm is divided into two stages: i) first mask creation and ii) wound and cell edge detection as summarized in Fig. 2. The first stage creates a mask with roughly defined wound edges, while the second stage refines the wound edges and detects cells within the wound. The algorithm description provided below is applicable with default parameters listed in Table 1.

**Figure 2:**
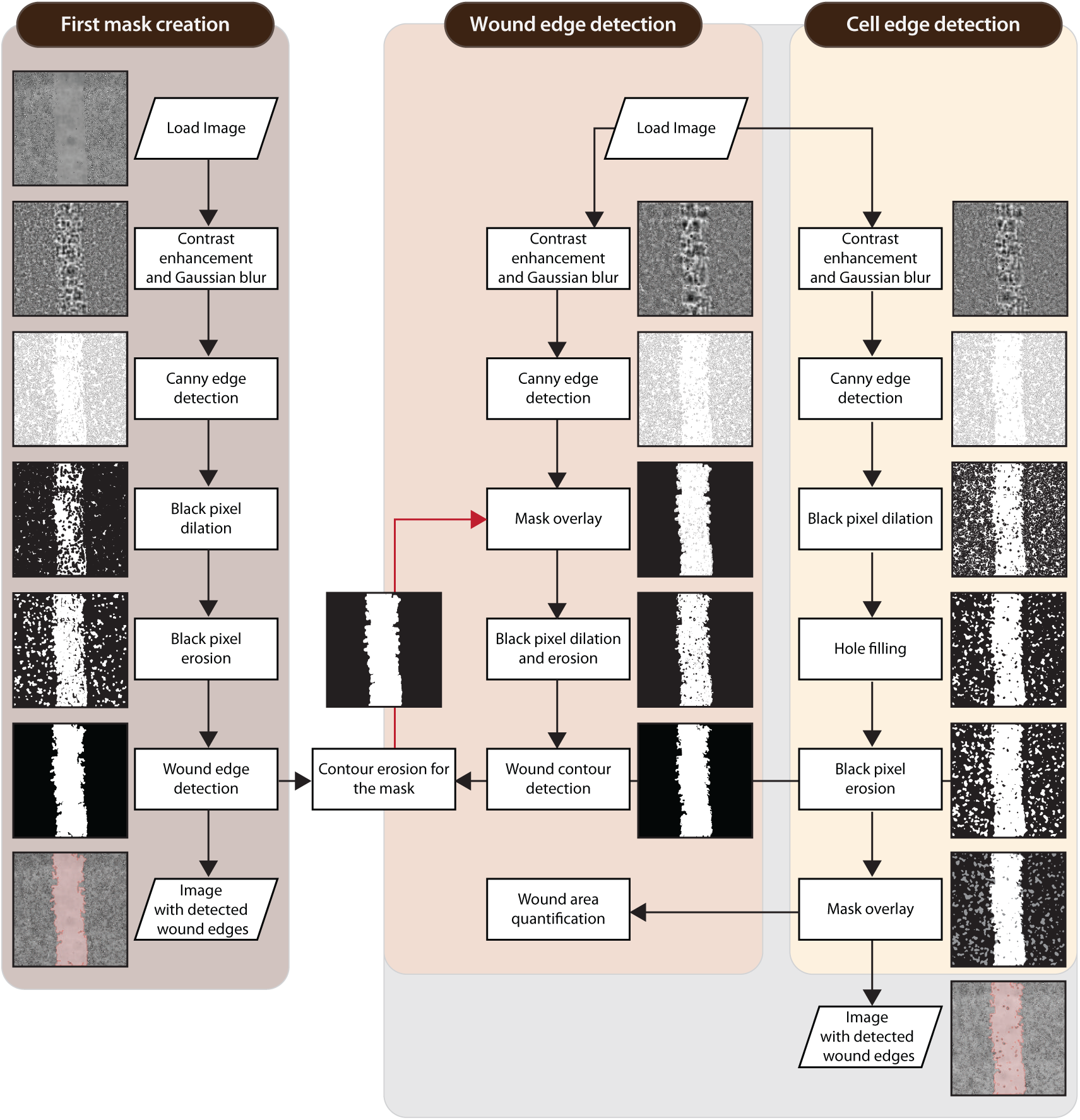
Schematic diagram of the CSMA algorithm workflow. Default parameters were used to obtain the representative images of the solvent control group of SW480-ADH colon cancer at every step.

**Table 1:** Default values applied for the adjustable parameters of the CSMA ImageJ plugin.

| Functional step in the pipeline | Parameter | Parameter setting value |  |  |
| --- | --- | --- | --- | --- |
|  |  | First mask creation | Wound edge detection | Cell edge detection |
| Contrast enhancement (CLAHE) | Contrast limit | 30 | 30 | 20 |
|  | Square grid size | 30 | 30 | 20 |
| Boundary dilation | Disk radius | 5 | 5 | 3 |
|  | Iterations | 2 | 1 | - |
| Boundary erosion | Disk radius | 5.5 | 5.5 | 3 |
|  | Iterations | 2 | 1 | - |
| Mask erosion | Disk radius | 5.5 | - | - |
|  | Iterations | 2 | - | - |
| Threshold | - | - | 0.15 | - |
| Cell filling | Kernel size | - | - | 25 |

First, the original image is preprocessed by applying contrast limited adaptive histogram equalization (CLAHE) image contrast enhancement method and Gaussian blurring. Gaussian blurring is achieved by convolving the input image with a Gaussian kernel of 9 *×* 9 pixels to remove noise signals. CLAHE is superior to simple histogram equalization since it generates fewer noise signals that may lead to artifacts in the processed image ^20^. After image preprocessing, wound edges are detected with the Canny edge detection method ^21^. The lower threshold value used to differentiate between candidate edges and true non-edges is calculated with Otsus method ^22^, while the higher threshold is fixed at 255.

Since the Canny method detects only individual cells within the cell monolayer on both sides of the wound, the next step is to fuse these individual cell boundaries into continuous wound front. This is done by first expanding the boundaries of the individual cells to merge them and then eroding them back using OpenCV morphological operations. Since users might have images of different sizes and resolutions, the parameters for all morphological operations are adjustable. The perimeter of the image is then padded with one row of black pixels to connect the front cells on two sides of the wound into a single continuous boundary. Lastly, the largest boundary (true wound boundary) filled with white pixels is redrawn on a black background with *cv2.drawCountours* function, creating a wound gap mask. Since morphological operations applied earlier distort the true edges of the wound, the redrawn wound boundary is eroded back to avoid overestimation in the output image.

The same operations are performed in the second stage of image processing. After the Canny edge detection method is applied in the wound edge detection pipeline, the black-and-white binary mask created earlier is overlaid on top of the newly obtained image to ensure that no residual holes remain in the cell monolayer in the output image. Morphological operations are applied as previously, but the parameter values are set lower for a more accurate representation of wound edges. After all boundaries, including true wound bondaries and individual cell boundaries, are detected on the black-and-white image with *cv2.findContours* function, the areas enclosed by these boundaries are calculated. If a boundary-enclosed area is smaller than 100 pixels or its ratio to the largest detected boundary is less than 0.15, it is classified as noise and discarded. The remaining boundaries are redrawn on a black background and filled with white pixels.

Similar steps are repeated to detect cells inside the wound. The output image with refined wound edges is obtained; however, the contrast limit and square grid size values for the CLAHE contrast enhancement function are reduced to minimize noise signals. Finally, the wound edge and cell edge detection pipelines are combined to produce an image with refined wound edges and cells within the wound.

At last, the number of white pixels representing the wound is calculated in the output image. The area of the gap is normalized and converted to the percentage as shown in Equation 1:

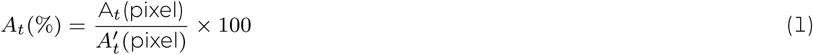

where *At* is the total cell-free area at a given time, and *A^′^*is the initial cell-free area.

The alternate width-based quantification method is also implemented using a similar image processing technique. As shown in Fig. S2, the difference is that the cell edge detection pipeline is removed from the algorithm, which significantly reduces the processing time, and the quantification of the gap width is performed by calculating the number of white pixels in every row. The mean of the gap width for every image is calculated as shown in Equation 2:

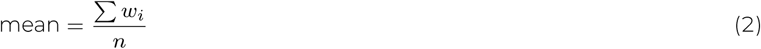

where *w_i_* is the gap width in *i^th^* row, and *n* is the number of rows.

The plugin produces three types of outputs: a .csv file with the calculated wound area or width for every time point in pixels and percentage formats, output images with detected wound boundaries, and a graph representing the closure of the wound over time.

The images acquired from treating SW480-ADH cells with DMSO, SVA, and WVA were analyzed by applying user-defined parameter values listed in Table 2.

**Table 2:** User-defined parameter values applied for the analysis of wound closure in SW480-ADH cells treated with DMSO, SVA, and WVA (user-adjusted values are in bold)

| Treatment<br>Replicate | DMSO |  |  |  | WVA |  |  |  | SVA |  |  |  |
| --- | --- | --- | --- | --- | --- | --- | --- | --- | --- | --- | --- | --- |
|  | 1 | 2 | 3 | 4 | 1 | 2 | 3 | 4 | 1 | 2 | 3 | 4 |
| First mask creation |  |  |  |  |  |  |  |  |  |  |  |  |
| Contrast limit | 5 | 5 | 10 | 1 | 10 | 10 | 5 | 10 | 5 | 10 | 5 | 5 |
| Square grid size | _____ 30 _____ |  |  |  | _____ 30 _____ |  |  |  | _____ 30 _____ |  |  |  |
| Dilation radius | 5 | 4 | 5 | 5 | 7 | 7 | 5 | 7 | 5 | 5 | 5 | 5 |
| Iterations | _____ 2 _____ |  |  |  | _____ 2 _____ |  |  |  | _____ 2 _____ |  |  |  |
| Erosion radius | _____ 5.5 _____ |  |  |  | _____ 5.5 _____ |  |  |  | _____ 5.5 _____ |  |  |  |
| Iterations | _____ 2 _____ |  |  |  | _____ 2 _____ |  |  |  | _____ 2 _____ |  |  |  |
| Mask erosion radius | 7 | 5 | 5 | 10 | 5.5 | 5.5 | 15 | 10 | 10 | 15 | 13 | 15 |
| Iterations | _____ 1 _____ |  |  |  | 2 | 2 | 3 | 3 | _____ 3 _____ |  |  |  |
| Wound edge detection |  |  |  |  |  |  |  |  |  |  |  |  |
| Contrast limit | 5 | 5 | 10 | 30 | 13 | 13 | 5 | 13 | 5 | 10 | 15 | 13 |
| Square grid size | _____ 30 _____ |  |  |  | _____ 30 _____ |  |  |  | _____ 30 _____ |  |  |  |
| Dilation radius | 4 | 4 | 5 | 6 | 7 | 7 | 8 | 7 | 6 | 7 | 9 | 5 |
| Erosion radius | 5.5 | 5.5 | 7.5 | 6.5 | 5.5 | 5.5 | 6 | 5.5 | 5.5 | 5.5 | 9.5 | 5.5 |
| Threshold | 0.15 | 0.15 | 0.15 | 0.1 | _____ 0.15 _____ |  |  |  | _____ 0.15 _____ |  |  |  |
| Cell edge detection |  |  |  |  |  |  |  |  |  |  |  |  |
| Contrast limit | 5 | 8 | 5 | 5 | 15 | 30 | 8 | 15 | 10 | 5 | 10 | 5 |
| Square grid size | _____ 30 _____ |  |  |  | 30 | 10 | 25 | 30 | 5 | 30 | 30 | 30 |
| Dilation radius | _____ 3 _____ |  |  |  | 3 | 3 | 4 | 3 | _____ 3 _____ |  |  |  |
| Erosion radius | _____ 3 _____ |  |  |  | 6 | 4.5 | 3 | 4.5 | _____ 3 _____ |  |  |  |
| Cell filling | 25 | 25 | 10 | 15 | 15 | 25 | 10 | 15 | 30 | 10 | 10 | 10 |

### Data analysis with MRI

Montpellier Ressources Imagerie (MRI) is a semi-automatic ImageJ tool for wound healing assay analysis ^23^. MRI tool relies on two different methods for image processing and requires the adjustment of four parameters. It can quantify the area of the wound on a time stack sequence of images and store it in a .csv file. Parameters in Table 3 were used for the analysis as indicated in their tutorial ^24^.

**Table 3:** Settings for the MRI ImageJ macro used for wound healing assay analysis.

| Parameter | Value |
| --- | --- |
| Method | Variance |
| Variance filter radius | 10 |
| Threshold | 50 |
| Radius open | 4 |
| Min. size | 10 000 |

### Data analysis with HTM

HTM Wound healing tool is a semi-automatic tool developed by Dr. Ivan Vorobyevs laboratory for the Scratch assay analysis. It is available in MatLab with no UI. Default parameters were used for the analysis. For demonstration purposes, the code was modified by changing the color and increasing the thickness of the detected boundaries in output images.

### Dynamics of wound closure

The dynamics of wound closure were initially assumed to be linear with a linear regression line fitted to all time points on the wound closure curve over 48 hours. The gradient or slope of this trendline was used to represent the wound closure rate, expressed as a percentage per hour.

Although linear dynamics of wound closure were observed for all detection methods, this pattern held true only for the early stages of wound closure. Careful observation revealed that the wound closure rate slowed down in the later stages (starting around 30 hours), deviating from the initial linear trend (see Fig. 1 and Fig. 4 for more details). Consequently, the dynamics of wound closure appeared to follow an exponential decay function rather than a linear curve. To better capture this behavior, a single exponential decay curve, *y* = *Ae^−λt^*, was used to estimate the dynamics of wound closure with *λ* representing the rate constant or decay constant. The comparison of its suitability against the linear model is shown in Fig. S4.

## RESULTS

### Evaluation of the CSMA plugin with default and user settings

Solvent-treated group of SW480-ADH cells was used to assess the accuracy of wound detection with the CSMA ImageJ plugin. The default parameters were tested against user-defined parameters, which were tuned according to the instructions provided for the plugin in the README file. As shown in Fig. 3, the number of cells found in the middle of the wound increased until they fused with the cells at the wound front after approximately 20 hours. The CSMA plugin successfully detected the cells in the middle of the gap at all time points, which allowed for more accurate area estimation. The final wound area was estimated to be 11.52% of the initial area (Fig. 3 and 4). The linear wound closure rate estimated for 48 hours was 1.83% per hour (*R*^2^ = 0.95). However, when two separate linear regressions were applied to two different regions of the graph with visually distinct wound closure rates, the following values were obtained: 2.48% per hour (*R*^2^ = 0.99) for the first 30 hours and 0.90% per hour (*R*^2^ = 0.90) for the remaining 18 hours (Fig. 4). The best-fitted exponential decay constant for the entire 48-hour period was 0.043 (*R*^2^ = 0.98).

**Figure 3:**
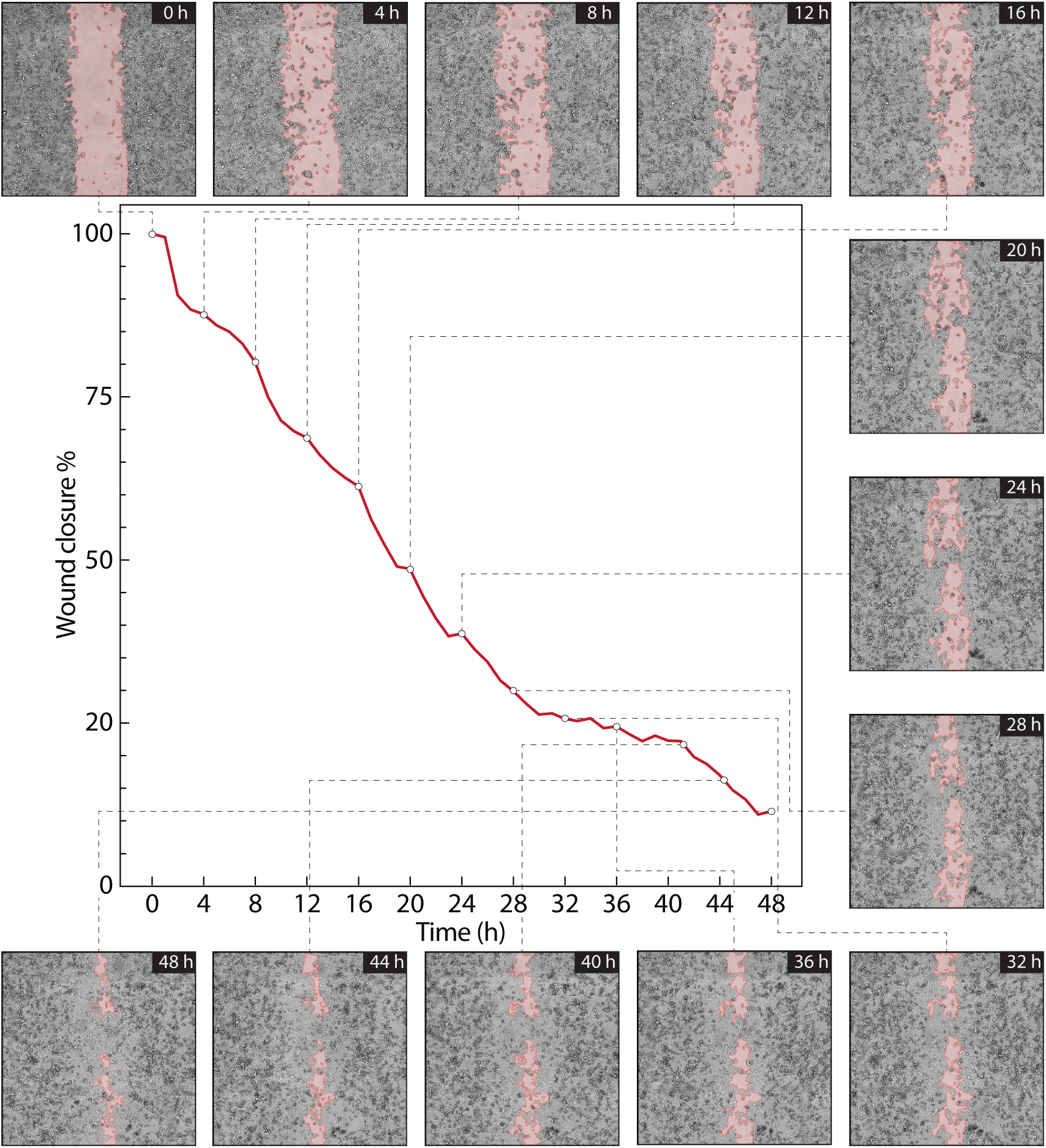
Wound closure in the vehicle control group of SW480-ADH colon cancer cells detected by the CSMA plugin with default settings. The graph representing the wound closure, which was generated automatically by CSMA algorithm and stored together with the output images.

**Figure 4:**
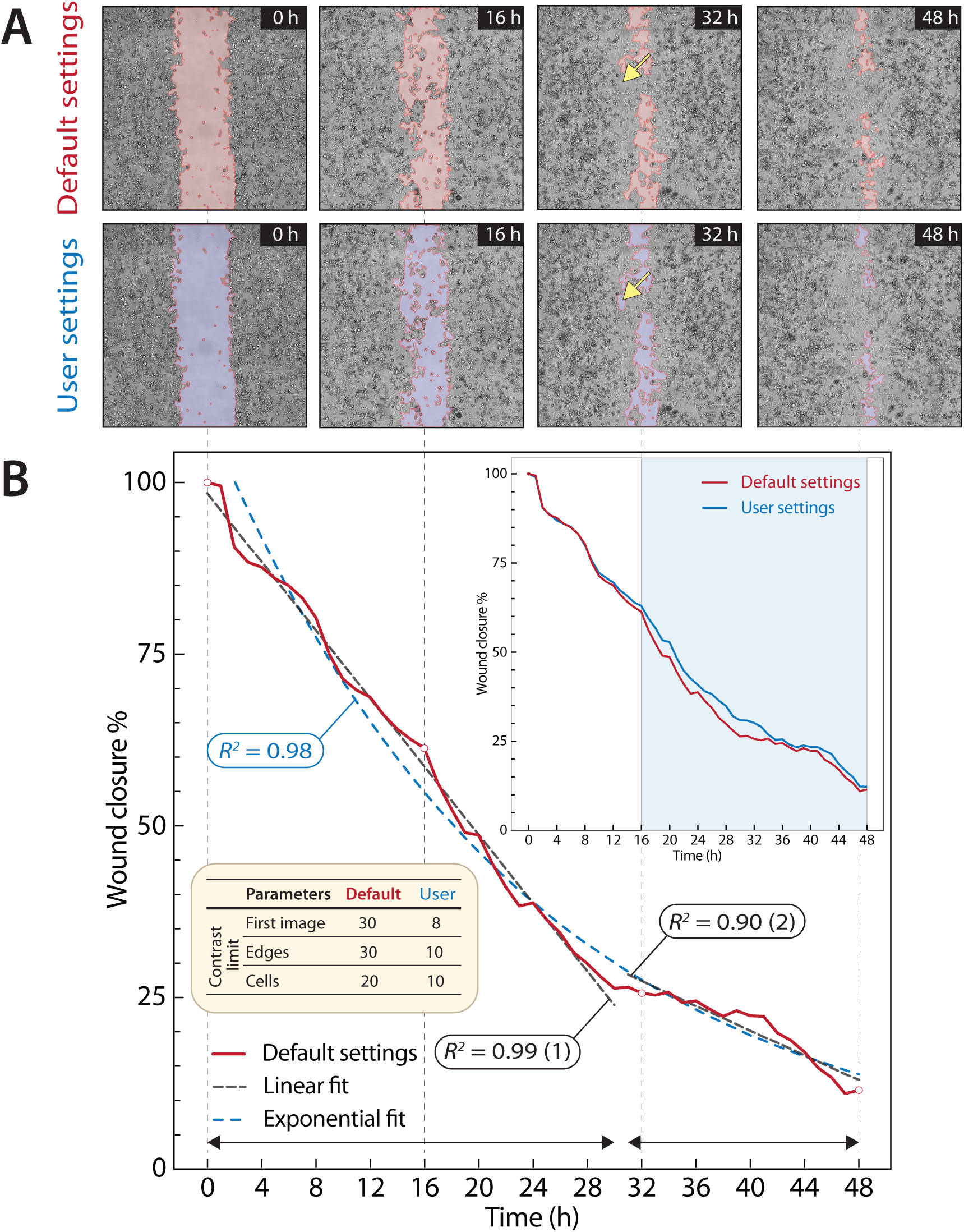
(a) Representative images of wound closure in the vehicle control group of SW480-ADH colon cancer cells detected by the CSMA plugin with default (red) and user-defined (blue) settings. The yellow arrow points to the overestimation of the cell-covered area for default settings. (b) Wound area closure vs. time. The inset shows the deviation of the curve between default and user setting from 16 hours on highlighted in blue area. The table within the graph displays the default and user-defined parameters used. The curve obtained for default setting has been fitted with linear and exponential functions demonstrating the superior performance of the exponential model (blue dashed line) over the linear (black dashed lines).

User manipulation of the plugin settings involved changing the contrast limits only since it was sufficient to achieve reliable quality of wound closure detection as demonstrated in Fig. 4. The final wound area as estimated with user-defined settings was 12.33%, which was slightly larger than with the default settings. The exponential decay rate reduced to 0.041 (*R*^2^ = 0.99).

During the first 16 hours, the wound area estimations obtained with default and user-defined parameters correlated closely since cells at the wound front moved uniformly, and cells in the middle of the wound remained separated from the wound front (Fig. 4). However, after 16 hours, the wound front came into proximity to the cells in the middle. The fusion of two wound fronts with cell-covered patches inside the wound led to an overestimation of the covered area for default settings. However, adjusting the contrast limit parameters minimized the area of regions falsely detected (Fig. 4A, yellow arrows).

### Performance comparison of CSMA, MRI, and HTM algorithms

To test the effectiveness of CSMA compared to other publicly available tools for wound healing assay, we selected MRI and HTM. To ensure a fair comparison between all the tools, we applied the recommended settings from the provided README file or source code for each one (Fig. 5). An additional dataset of 769-P cells was chosen to ensure that our plugin worked properly on images with different characteristics (Supplementary Fig. S3).

**Figure 5:**
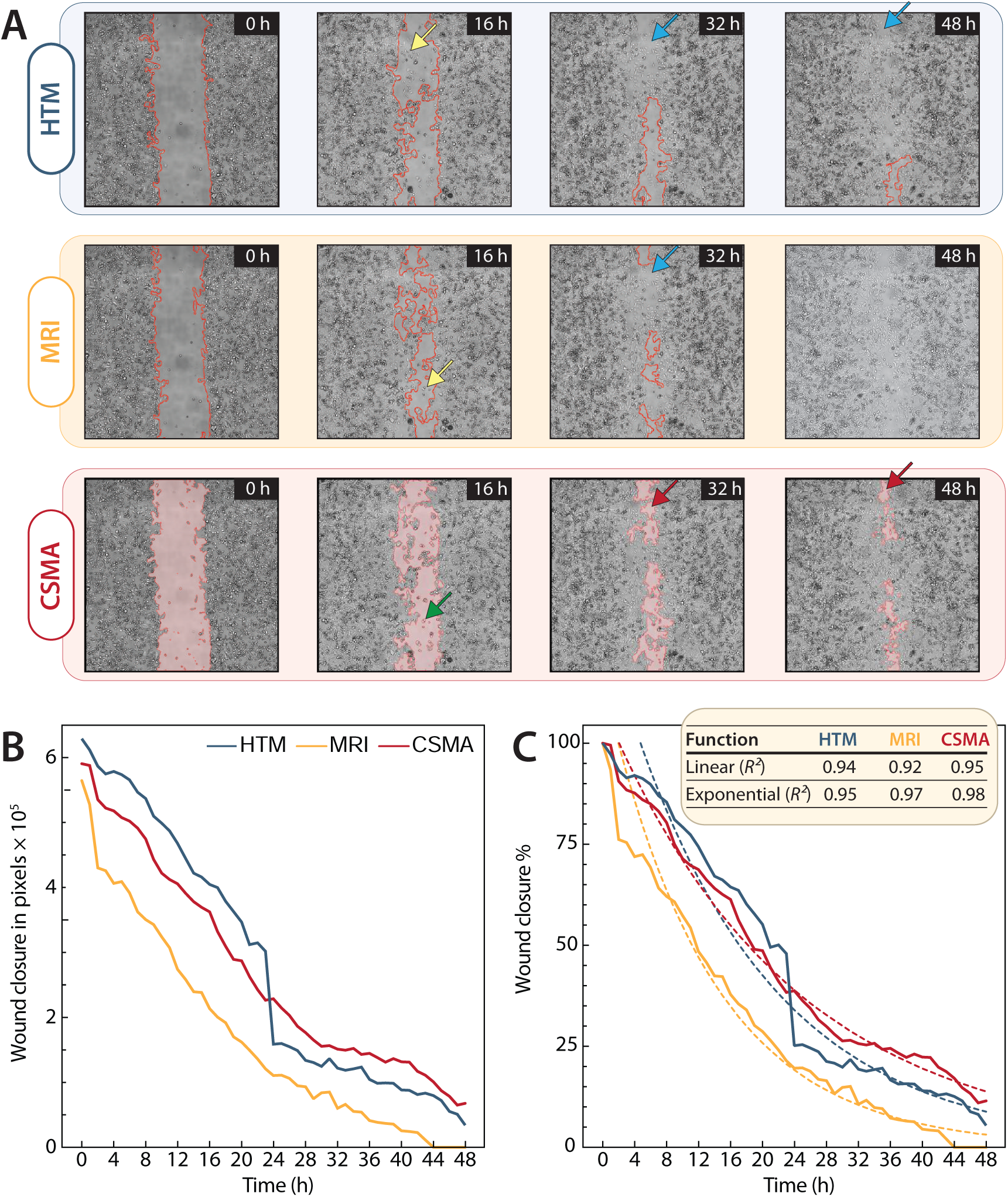
(a) Representative images of wound closure in the vehicle control group of SW480-ADH colon cancer cells as detected by HTM MATLAB code, the MRI ImageJ macro, and the CSMA ImageJ plugin. Default settings were used for all three wound detection tools. (b) Quantification of the wound area closure over 48 hours expressed in pixels. (c) Quantification of the wound area closure over 48 hours expressed as a percentage. The table displays the *R*^2^ values for both linear and exponential fits demonstrating the superior performance of the exponential function over linear.

Fig. 5A highlights differences in how cell boundaries were detected by different algorithms. While both HTM and MRI had some challenges in detecting cells at the center of the wound area (yellow arrows), CSMA detected these cells with high precision (green arrow). Further, HTM and MRI could not detect the empty regions in the middle of the wound after 32 hours (blue arrows), leading to biased wound area estimations. In contrast, our plugin accurately detected these regions (red arrows). CSMA showed a smaller initial wound area compared to HTM because it successfully detected the cells inside the wound (Fig. 5B). On the other hand, MRI demonstrated a smaller initial wound area than CSMA presumably because of the overestimation of the wound front (see Fig. 5A). Finally, CSMA produced a much smoother wound closure curve than HTM, which falsely detected only the largest remaining wound region as the true wound at approximately 24 hours neglecting the smaller unoccupied areas (Fig. 5B and C).

CSMA also effectively detected wound boundaries and cells within the wound in the 769-P dataset (Supplementary Fig. S3), accurately tracking the migration of cells within the wound. The wound area estimations of CSMA and HTM correlated very closely, except for the sudden drop at 22 hours detected by HTM, which was caused by the disconnected detection of the wound area while only the largest area was used for the estimation. MRI underestimated the area of the wound at all times most probably due to the specificities of the detection algorithm.

In general, CSMA accurately detected cells within the wound in three replicates of SW480-ADH colon cancer cells treated with DMSO. The mean *R*^2^ values obtained from three replicates for the exponential best-fit curves for CSMA, HTM, and MRI were 0.97*±*0.01%, 0.95*±*0.03%, and 0.91*±*0.08%, respectively (see Supplementary Fig. S4). As a result, the wound closure curve of CSMA fitted the exponential model better than that of HTM and MRI as demonstrated by the calculated *R*^2^ values.

### Effect of SVA and WVA on the migration of SW480-ADH cells

The effect of SVA and WVA on the migration of human colon cancer cells was assessed by the CSMA plugin (Fig. 6A). SW480-ADH cells treated with 100*µ*M WVA demonstrated the lowest mean linear wound closure rate of 0.78*±*0.22 % per hour (*R*^2^ = 0.93*±*0.05), compared to 1.17*±*0.30 % per hour (*R*^2^ = 0.98*±*0.01) in the group treated with 10 nM SVA, and 1.63*±*0.21 % per hour (*R*^2^ = 0.96*±*0.02) in the solvent control group. The mean exponential decay constants obtained from four replicates were estimated as follows: 1.03*×*10*^−^*^2^ *±*3.30*×*10*^−^*^3^ (*R*^2^ = 0.93 *±* 0.06) for WVA, 1.78 *×* 10*^−^*^2^ *±* 8.18 *×* 10*^−^*^3^ (*R*^2^ = 0.97 *±* 0.01) for SVA, and 2.83*×*10*^−^*^2^ *±* 6.99 *×* 10*^−^*^3^ (*R*^2^ = 0.96 *±* 0.02) for the solvent control group. The average final wound areas were 27.22 *±* 8.25 %, 45.20 *±* 12.74 %, and 59.89 *±* 8.22 % for the DMSO, SVA, and WVA groups, respectively (Fig. 6B).

**Figure 6:**
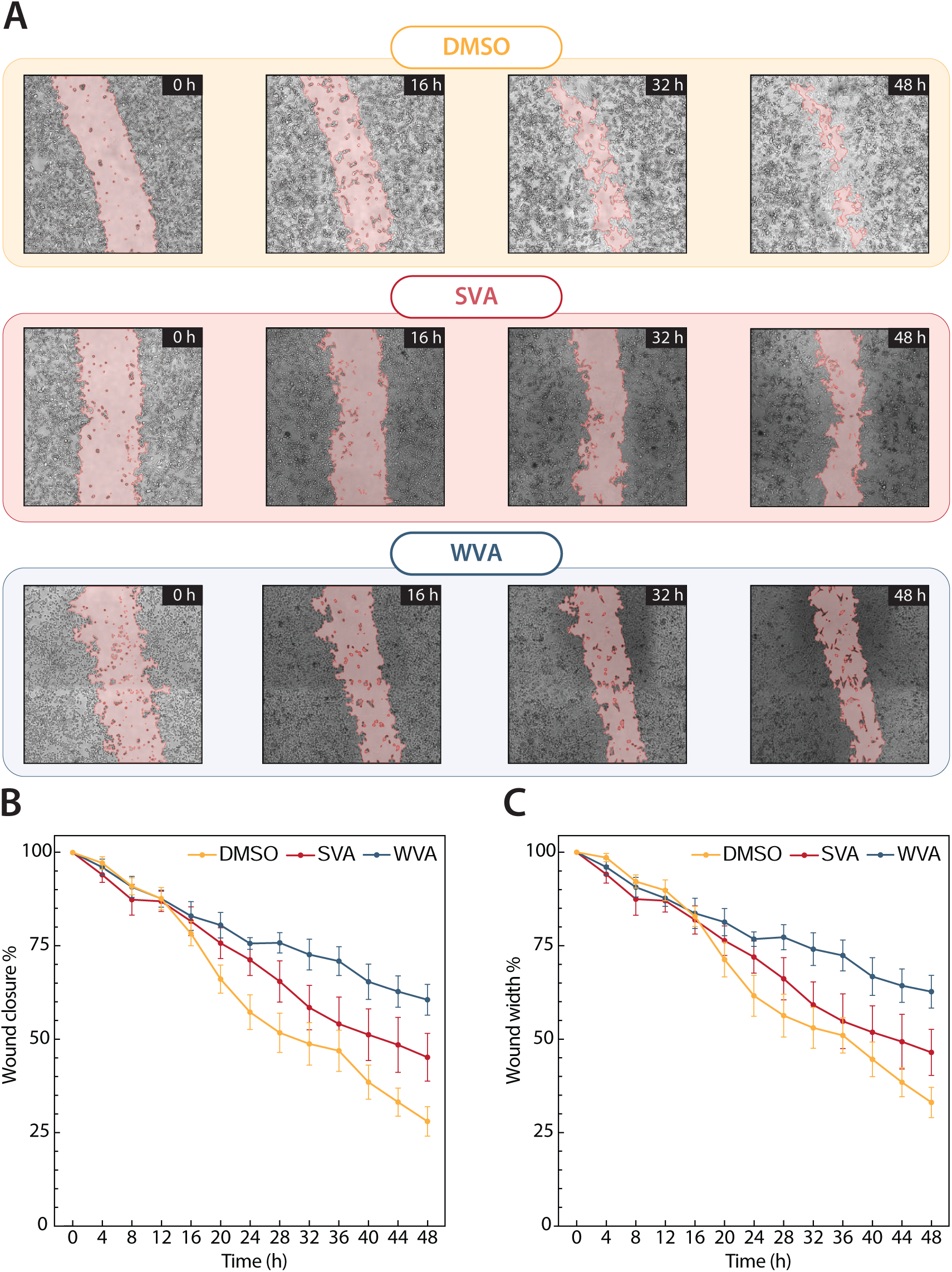
(a) Representative images of wound closure in SW480-ADH colon cancer cells treated with DMSO, 10nM SVA, and 100*µ*M WVA detected by the CSMA ImageJ plugin with user-defined parameters listed in Table 2. (b) Quantification of the mean wound area closure over 48 hours obtained from four replicates. (c) Quantification of the mean wound width over 48 hours obtained from four replicates.

## DISCUSSION

The CSMA plugin demonstrated high-precision edge detection in all tested datasets by detecting migrating cells in the middle of the wound and minimizing the overestimation of wound edge boundaries.

The performance of the plugin with default and user-defined parameters was comparable, and the final wound area estimations for both vehicle control group SW480-ADH and 769-P cells were less than 1% difference, demonstrating high compatibility of the default parameters with different imaging conditions. However, with user-defined parameters, the wound closure was more accurate in the vehicle control group SW480-ADH since the false detection of empty regions was minimized. Importantly, such improvement in wound detection resulted from the adjustment of only the contrast limits, while the rest of the parameters remained the same. In all analyzed datasets, the user-defined contrast limits of wound edge detection and first mask contrast limit values matched closely, which is why keeping these two parameters the same is a good starting point for fine-tuning the plugin settings as listed in Tables 1 and 2.

The detection of cells in the middle of the wound was precise for both the SW480-ADH and 769-P datasets. For instance, the plugin accurately detected the changing boundaries of a migrating cell front in the 769-P dataset with default settings (Supplementary Fig. S3). This suggests that the default parameters of the CSMA plugin may serve as a universal starting point for accurate wound detection across various imaging conditions. Other detection tools were unable to account for these cells in their calculations, suggesting that our plugin is one of the few freely available options that offer such high precision in wound area closure detection. Additionally, other tools failed to detect empty regions occasionally, leading to inaccurate wound area detection. The CSMA plugin also outperforms others in terms of image processing speed. For example, while HTM takes about 4.19 *×* 10*^−^*^1^ *±* 6.51 *×* 10*^−^*^3^ seconds to process an image, CSMA requires only 2.26 *×* 10*^−^*^1^ *±* 5.49 *×* 10*^−^*^4^ second with default settings.

The datasets used for testing the effect of various VDR agonists on SW480-ADH migration were more difficult to analyze for several reasons. First, uneven and constantly changing illumination impeded edge detection because the borders of dark and light regions were detected as the wound edges by the Canny method, which is based on detecting the regions of sharp illumination changes ^21^. This drawback was minimized by carefully adjusting the contrast limit and square grid size. Increasing the contrast limit and decreasing the square grid size improved the detection of smaller elements, such as single cells, while decreasing the contrast limit and increasing the square grid size helped to reduce the detection of false edges.

In addition, it is occasionally possible for the “field of view” and the “location of the wound in the image” to be slightly shifted as a result of the image stitching process or a minor disturbance of the imaging device inside the incubator during the image acquisition procedure. Since the CSMA plugin applies the mask from the previous image to the current one, this shift resulted in an inaccurate wound detection with the default parameters. However, this limitation was overcome by increasing the mask erosion parameter. It is advised to accompany such adjustments with increasing the dilation radius to maintain accurate detection of the true wound edges. Lastly, the microscope’s focus likely shifted over time, causing cells inside and at the front of the wound to appear larger and blurrier in later images. Although the resulting sudden drops in the wound area could not be entirely corrected, the plugin still accurately detected small elements and wound edges.

Precise detection of wound edges and cells within the wound allowed CSMA to achieve high-accuracy area detection and produce smooth wound closure curves. As a consequence, wound closure curves generated by CSMA fit the linear and exponential models better than those generated by HTM and MRI (Supplementary Fig. S4). Although both the linear and exponential regression curves have *R*^2^ values above 0.90, the exponential regression curve is more suitable due to its higher *R*^2^ value. This is because exponential curves can accurately fit both visually linear and exponential wound closure patterns, whereas the linear fit aligns well only in cases of linear closure. For example, as demonstrated in Fig. 4 and 5, a single linear fit line cannot adequately describe the cell behavior over the entire period since different time intervals exhibit varying closure speeds. Therefore, two linear trendlines are required to capture the true behavior of this curve, which is both time-consuming and potentially misleading. In contrast, a single exponential decay curve fits the data well and allows the behavior to be described with only one decay rate.

One limitation of the study is the inconsistency in the initial wound width due to variations in the force applied to the pipette tip. Additional challenges include low image quality, cells with suboptimal adhesion properties such as being overly adhesive or non-adhesive, number of adjustable parameters, and potential installation issues due to operating system differences and unique user settings.

Though CSMA offers two modes of wound closure quantification, we found that the area quantification method was more accurate than width because it accounted for cells proliferating in the middle of the gap.

## CONCLUSION

Our study demonstrates that the newly developed CSMA ImageJ plugin offers a significant advancement in the accurate quantification of wound closure in cell migration assays. By addressing the limitations of existing methods, which often fail to detect cells within the wound gap and result in biased measurements, our plugin provides a more reliable and precise approach for evaluating cell migration and wound healing. The ability of our tool to identify empty regions and avoid overestimation of the cell-covered area further validates its effectiveness. Additionally, accurately detecting wound closure at all time points allows for the correct estimation of wound closure dynamics such as its rate. This innovative solution not only enhances the accuracy of wound closure assessments but also offers a user-friendly platform for researchers to obtain robust and reproducible results in cell migration studies such as in studying the metastatic properties of cancer cell lines.

## ACKNOWLEDGMENTS

We extend our sincere gratitude to Daniyar Kakimbekov from the Department of Computer Science at Nazarbayev University for his invaluable advice and support throughout this project.

## SUPPLEMENTARY DATA

**Figure S1:**
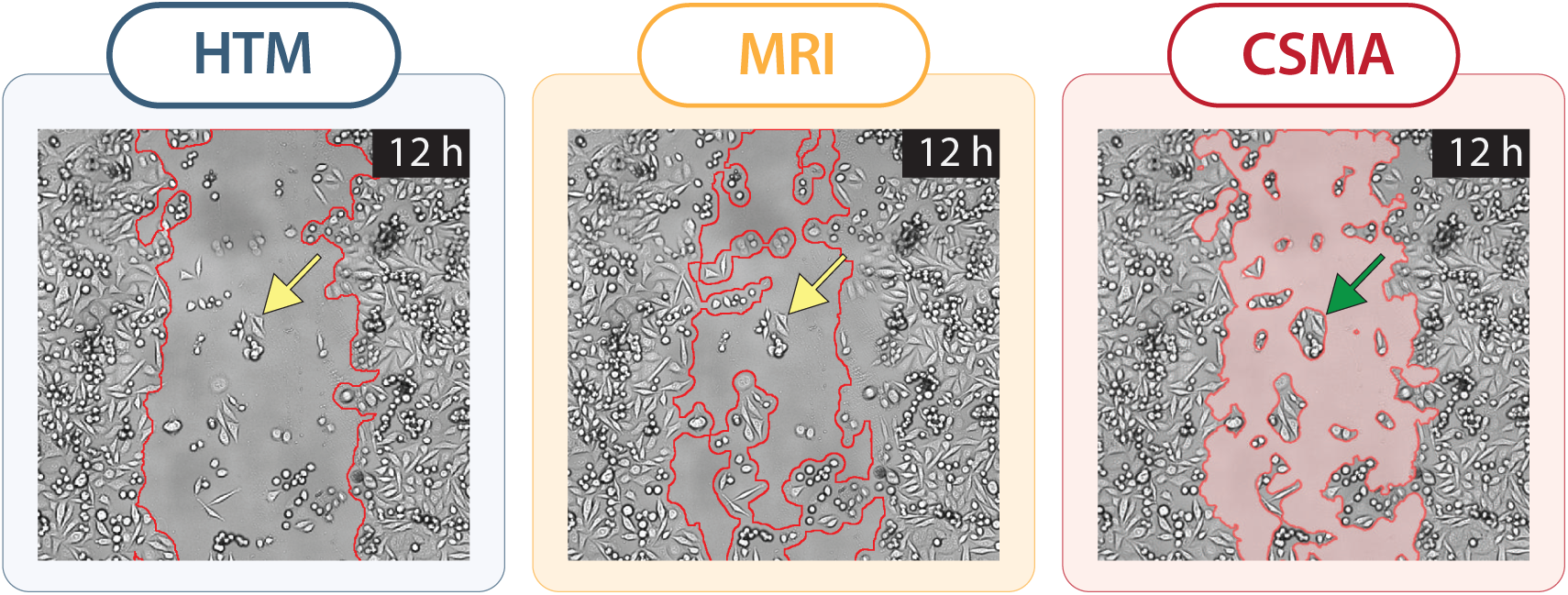
Representative images of wound closure at 12 hours in SW480-ADH colon cancer cells treated with DMSO and detected using the CSMA ImageJ plugin, MRI ImageJ macro, and HTM MatLab code with default settings. The two yellow arrow indicate the misdetection of cells inside the would for HTM and MRI compare proper detection via CSMA (green arrow).

**Figure S2:**
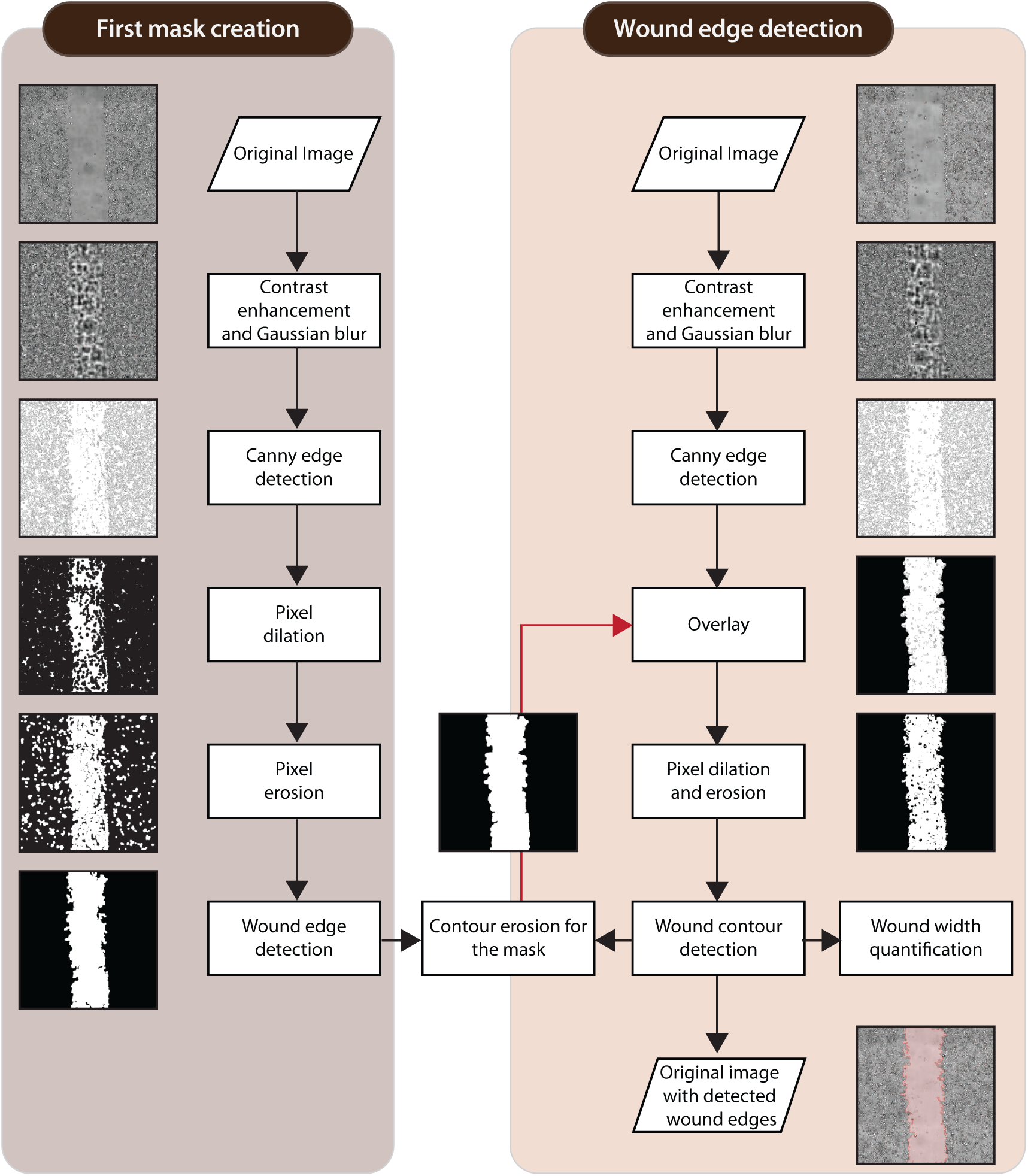
Schematic diagram of the CSMA algorithm workflow for wound width detection and quantification. Default parameters were used to obtain the representative images for every step.

**Figure S3:**
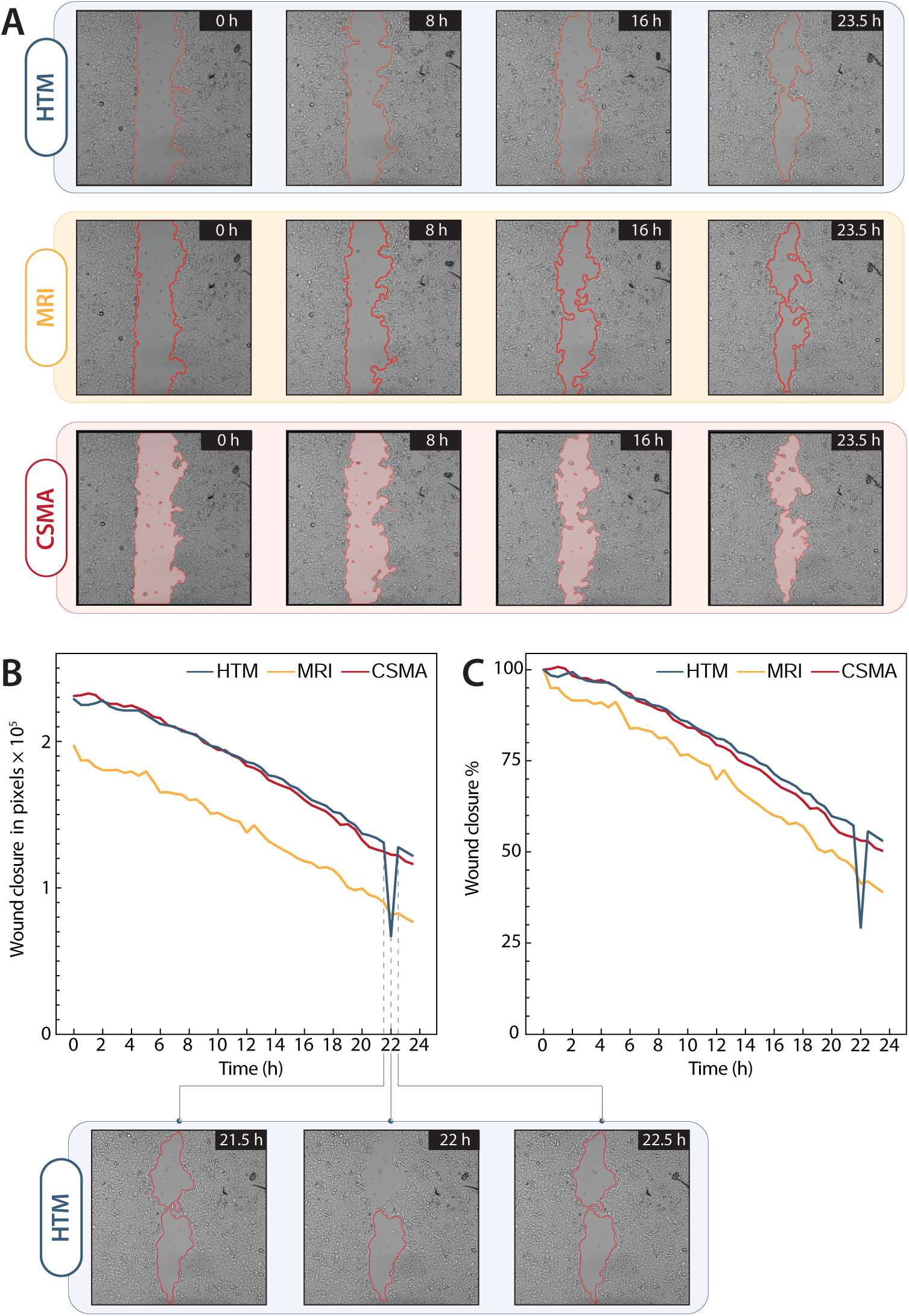
Representative images of wound closure in mutant 769-P renal cancer cells detected using the CSMA ImageJ plugin, MRI ImageJ macro, and HTM MATLAB code with default settings. (b) Quantification of wound area closure in pixels over 23.5 hours. (c) Quantification of wound area closure as percentage over 23.5 hours. The drop in wound closure area corresponds to a smaller area detected at 22 hours, as shown in the images below the graph alongside the 21.5- and 22.5-hour time points.

**Figure S4:**
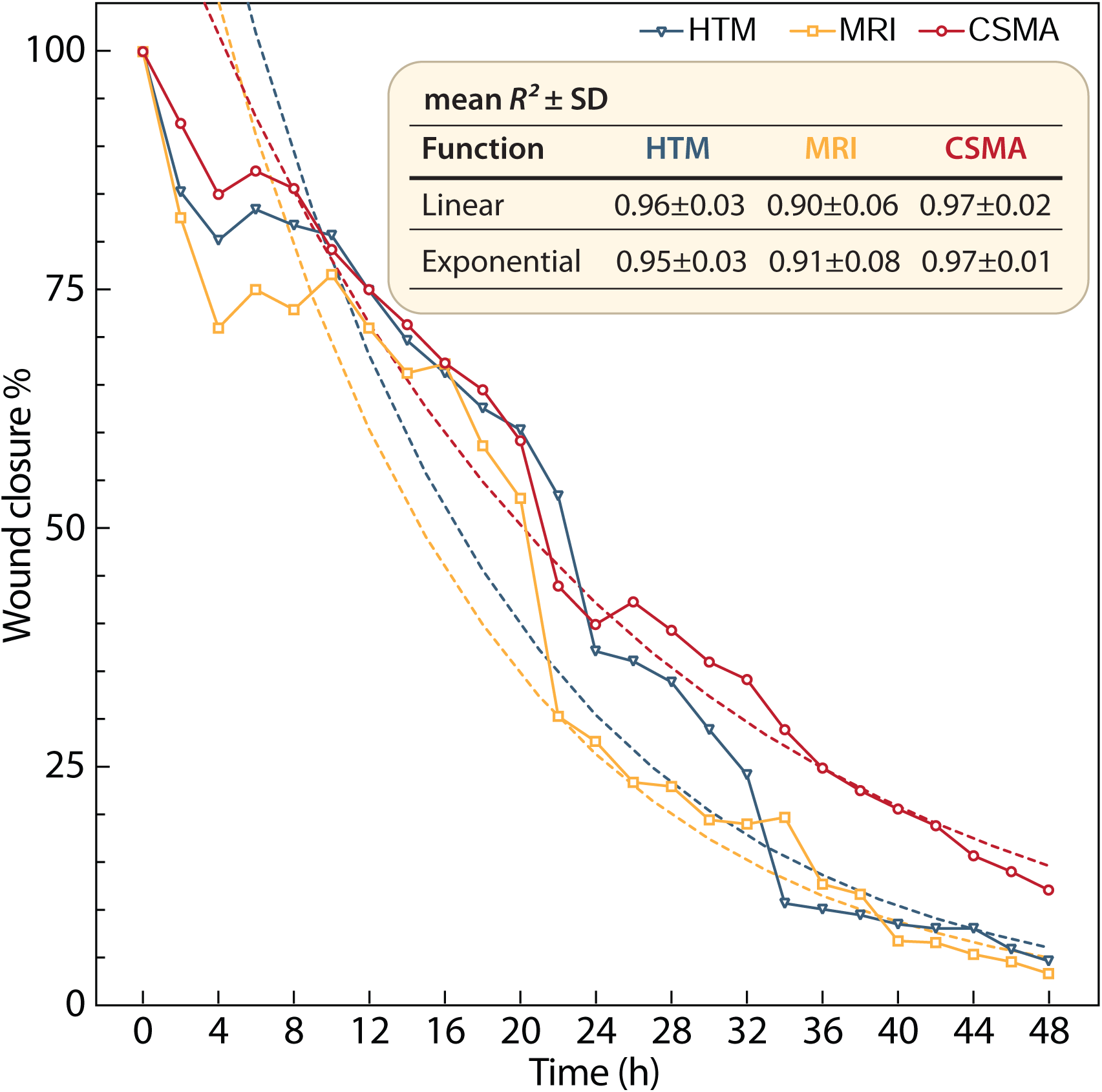
Representative graphs of the wound area closure vs time for the vehicle control group of SW480-ADH colon cancer cells detected by HTM, MRI, and CSMA algorithms. Both linear and exponential curves were fitted to determine the *R*^2^ values. The table within the graph displays the mean *R*^2^ for both linear and exponential regression curves obtained from three independent measurements.

